# Amniotic fluid stem cell extracellular vesicles promote lung development via TGF-beta modulation in a fetal rat model of oligohydramnios

**DOI:** 10.1101/2024.06.18.599591

**Authors:** Fabian Doktor, Rebeca Lopes Figueira, Victoria Fortuna, George Biouss, Kaya Stasiewicz, Mikal Obed, Kasra Khalaj, Lina Antounians, Augusto Zani

## Abstract

Oligohydramnios (decreased amniotic fluid volume for gestational age) is a severe condition associated with high morbidity and mortality mainly due to fetal pulmonary hypoplasia. Currently, there are limited treatment options to promote fetal lung development. Administration of stem cells and their derivates have shown promising regenerative properties for several fetal and neonatal diseases related to arrested lung development. Herein, we first characterized pulmonary hypoplasia secondary to oligohydramnios in a surgical rat model. Experimental induction of oligohydramnios led to impaired lung growth, branching morphogenesis (fewer airspaces with decreased *Fgf10*, *Nrp1*, *Ctnnb1* expression), proximal/distal progenitor cell patterning (decreased Sox2 and Sox9 expression), and TGF-β signaling. We then tested antenatal administration of extracellular vesicles derived from amniotic fluid stem cells (AFSC-EVs). In oligohydramnios lungs, AFSC-EV administration improved lung branching morphogenesis and airway progenitor cell patterning at least in part through the release of miR-93-5p. Our experiments suggest that AFSC-EV miR-93-5p blocked SMAD 7, resulting in upregulation of pSMAD2/3 and restoration of TGF-β signaling. Conversely, oligohydramnios lungs treated with antagomir 93-5p transfected AFSC-EVs had decreased branching morphogenesis and TGF-β signaling. This is the first study reporting that antenatal administration of stem cell derivatives could be a potential therapy to rescue lung development in fetuses with oligohydramnios.

**Highlights:**

- Pulmonary hypoplasia secondary to oligohydramnios in fetal rats is characterized by impaired TGF-β signaling.
- AFSC-EV administration improves fetal lung branching morphogenesis and airway progenitor cell patterning.
- AFSC-EV effects are mediated at least in part via modulation of TGF-β signaling by the release of miR-93-5p from AFSC-EVs.

## Introduction

About 3-10% of pregnant women develop a complication called preterm premature rupture of the membranes (PPROM).^1–3^ This is defined as rupture of fetal chorioamniotic membranes before onset of labor at less than 37 weeks of gestation and is responsible for about a third of preterm deliveries.^3,4^ Babies born from pregnancies affected by PPROM have pulmonary hypoplasia, a condition characterized by incomplete lung development *in utero*.^5^ The main cause of fetal pulmonary hypoplasia is the leakage of amniotic fluid through the ruptured membranes resulting in a decreased volume for gestational age, also known as oligohydramnios.^6^ In these fetuses, oligohydramnios is the main determinant for severe neonatal respiratory morbidity and mortality.^7^ Attempts have been made to replenish the normal amniotic fluid volume by serial or continuous infusions with non-amniotic fluids during the mid- and third trimester.^8–11^ However, randomized controlled trials have shown little or no benefit with amnioinfusion for perinatal survival, rather they have highlighted possible associated complications.^8,10,11^ One could speculate that re-establishing just fluid volume is not enough, as the amniotic fluid is highly enriched with nutrients that are required for normal organogenesis.^12^ In particular, lung development is a complex process regulated by multiple signaling pathways involving several molecular species, such as miRNAs, proteins, and lipids.^13–15^ Several studies have shown that some miRNAs are missing or dysregulated in human and animal fetal hypoplastic lungs and that a therapy that supplements the several missing miRNAs and bioactive molecules involved in lung development could be a strategy to rescue pulmonary hypoplasia.^13–25^ Extracellular vesicles (EVs) are ideal candidates to deliver a complement of several biomolecules, as they are lipid-bound nanoparticles that carry miRNAs, proteins, and lipids as cargo, which they transfer to target cells to induce biological responses.^26^ For this reason, EVs have been considered as a multi-targeted therapy for conditions where a single drug treatment would not suffice as multiple molecules and/or pathways are missing or dysregulated, such as bronchopulmonary dysplasia.^27^ A potential novel antenatal therapy for oligohydramnios-induced pulmonary hypoplasia could similarly be based on a stem-cell based regenerative therapy. In particular, stem cells derived from the amniotic fluid (AFSC) could be an ideal EV source as they differentiate into epithelial lung lineages, reduce lung fibrosis, and repair alveolar epithelial injury.^28–30^ AFSC-EVs have shown promise in models of fetal pulmonary hypoplasia secondary to congenital diaphragmatic hernia (CDH).^13,31,32^ In experimental CDH, antenatal administration of AFSC-EVs resulted in the rescue of branching morphogenesis and improvement of epithelial and mesenchymal differentiation by restoring dysregulated signaling pathways that are relevant to lung development.^13,33^ These beneficial effects were exerted via the release of the EV RNA cargo, which is enriched in miRNAs that are critical for branching morphogenesis, such as miR 17∼92 cluster and its paralogues.^13,16,25,34,35^ Enzymatically digesting the AFSC-EV RNA cargo or blocking specific miRNAs regulating lung developmental processes with antagomirs did not reproduce the beneficial effects observed in hypoplastic lungs upon AFSC-EV administration.^13^

In this study, we used an experimental model of oligohydramnios-induced pulmonary hypoplasia in fetal rats and investigated the dysregulated signaling pathways that are affected. We discovered that fetal hypoplastic lungs have impaired branching morphogenesis with dysregulation of TFG-β/SMAD signaling. Moreover, we evaluated the effects of antenatal administration of rat AFSC-EVs, which resulted in restoration of branching morphogenesis and improvement of lung progenitor cell differentiation. Using antagomirs, we conducted mechanistic studies that revealed that these beneficial effects on lung development are at least in part exerted via the release of miR-93-5p contained in the AFSC-EV cargo.

## Material and Methods

### Extracellular Vesicle isolation, characterization and tracking

C-kit+ rat AFSCs were isolated at E12.5 and cultured to 80% confluence in alpha-minimal Essential Media (αMEM) supplemented with 15% fetal bovine serum (FBS) as well as 20% Chang medium. Subsequently, AFSCs were grown in 7.5% exosome-depleted FBS (18 hours) to obtain conditioned medium. 300g and 1200g differential centrifugation steps were employed to remove dead cells and debris. Before ultracentrifugation at 100,000g (14 hours, SW 32Ti Beckman Coulter), the cleared medium was filtered (0.22μm filter).^36,37^

According to the International Society for Extracellular Vesicles (ISEV) guidelines, AFSC-EVs were characterized for size via nanoparticle tracking analysis, for morphology via transmission electron microscopy, and for the expression of canonical EV-markers via Western blot.^38,39^

For nanoparticle tracking analysis via NTA 3.4 Build with sCMOS camera, 5x 40s were captured at 23°C. For transmission electron microscopy, AFSC-EVs underwent fixation in 4% paraformaldehyde (PFA), placed onto a charged formvar-carbon coated copper grid, contrasted via uranyl-oxalate, embedded in methyl cellulose-UA, and then imaged from 25 kx to 100 kx magnification via Tecnai 20 (Thermo Fisher Scientific). For expression of canonical EV protein markers, AFSC-EV protein was extracted via a phosphatase and protease inhibitor (Roche) containing tissue extraction buffer (Thermo Fisher Scientific). Protein concentration was then measured with a Pierce Bradford assay (ThermoFisher Scientific) and analyzed for the expression of cell differentiation marker 63 (CD63), Tumor susceptibility 101 (TSG101), and Flotillin-1 (Flot1). Cellular lysate was used as a positive control for Calnexin (CANX).

AFSC-EV RNA cargo was labeled via Exo-GLOW™ Exosome Labeling kit (SBI System Biosciences) according to the manufacturer’s protocol. The Exo-GLOW™ labeled AFSC-EVs were added to the medium of fetal hypoplastic lung explants 1h prior to imaging, co-incubated with DAPI and imaged via 2-photon Leica SP8 microscope. Unlabeled AFSC-EVs were used as a negative control.

### Experimental model of pulmonary hypoplasia secondary to oligohydramnios

All animal experiments were approved by the Animal Care Committee (Animal use protocol #49892 and #65210) of The Hospital for Sick Children, Toronto, Canada according to the Canadian Council on Animal Care Guidelines.

Pregnant Sprague-Dawley rats were anesthetized and a midline laparotomy was performed to expose the uterine horns at E16.5. The amniotic sac was punctured largely enough to allow subsequent continuous loss of amniotic fluid without closure. Control dams underwent laparotomy alone. On E19.5, dams were euthanized, fetuses were delivered via C-section and their lungs were removed. To limit variability, we used only oligohydramnios lungs with a lung to body weight ratio <3%. Lungs were grown *ex vivo* as lung explant cultures for 24 hours on nanopore membranes (Thermo Fisher Scientific) supplemented with culture medium alone (Dulbecco’s Modified Eagle Medium+10% fetal bovine serum+0.5% penicillin/streptomycin) or rat AFSC-EVs (0.5%, v/v). The left and right lung lobes were assigned in a random fashion to either fixation with 10% para-formaldehyde (PFA) for 24h or snap frozen and stored at -80°C.

### Histology and pulmonary hypoplasia assessment

Fixed fetal rat lungs were embedded in paraffin or OCT (Sakura Finetek). The formalin-fixed-paraffin-embedded (FFPE) blocks were sectioned into 5μm thick slides and OCT embedded lung tissue was sectioned into 12 μm thick slides. For histological assessment, lungs were stained with hematoxylin and eosin (H&E). After deparaffinization, slides were stained first with hematoxylin for approximately 10min and with eosin for approximately 2min. According to the recommendation of the American Thoracic Society, severity of pulmonary hypoplasia was assessed by determining the radial airspace count (RAC) and the mean linear intercept (MLI).^40^ RAC counting was performed blinded and MLI was assessed via a semi-automated quantification method published previously.^41^ The lung-to-body weight ratio was also used to assess the severity of pulmonary hypoplasia.

### RNA extraction, cDNA synthesis and RT-qPCR

Lung specimens were stored at -80°C. TRIzol™ Reagent (ThermoFisher Scientific) was used according to the manufacturer suggested protocols to isolate RNA. A NanoDrop™ spectrophotometer was utilized to quantify concentration and purity of the isolated RNA. Subsequent cDNA synthesis was performed (Invitrogen™ SuperScript™ VILO™ cDNA Synthesis Kit) and cDNA was stored at -20°C. RT-qPCR experiments were performed according to the MIQE guidelines^42^ with 5.5μL cDNA, 5.5μL SYBR™ Green Master Mix (Applied Biosystems™, ThermoFisher Scientific, SYBR™ Green PCR Master Mix) and 0.25μL forward- and reverse primer each. Primers are listed in **Table 1**. Briefly, lung branching morphogenesis was assessed via gene expression of fibroblast growth factor 10 (*Fgf10*), neuropilin-1 (*Nrp1*) and β- catenin (*Ctnnb1*). Lung progenitor cell expression was assessed via SRY-box transcription factor 2 and 9 (*Sox2* and *Sox9*). Smooth muscle cell gene expression was quantified via actin alpha 2 (*Acta2)* and transforming growth factor beta 1,2 and 3 as well as transforming growth factor beta receptor 1 and 2 (*Tgfb1, Tgfb2, Tgfb3, Tgfbr1, Tgfbr2*) gene expression was quantified to analyze the TGF-β signaling pathway. In total, 45 cycles (denaturation: 95 °C, annealing: 58 °C, extension: 72 °C) were performed on a ViiA7 qPCR system. Gene expression was calculated using the 2^ΔΔ^CT method and normalized to glyceraldehyde-3-phosphate dehydrogenase (*Gapdh*), as *Gapdh* was expressed equally across the experimental groups.

**Table 1:**
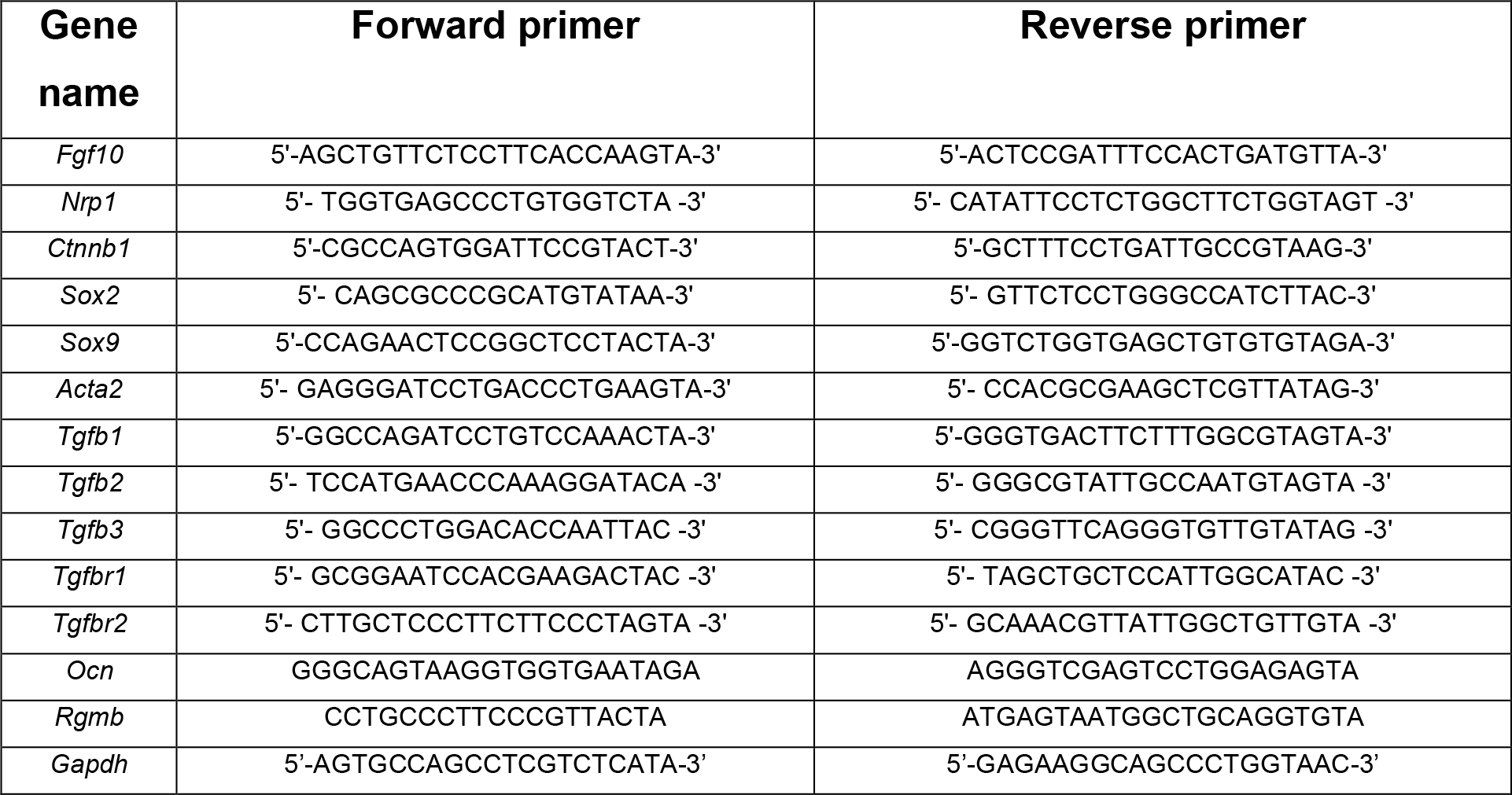
Primer sequences used for RT-qPCR.

### Immunofluorescence experiments

FFPE fetal lung rat tissue was deparaffinized, antigen retrieval was achieved by immersing samples in citrate buffer under high pressure, and following incubation with 3% bovine serum albumin (BSA) sections were incubated with primary antibodies overnight (**Table 2**) with subsequent 1h secondary antibody incubation. The slides were stained with 4′,6-diamidino-2-phenylindole (DAPI) (Thermo Fisher Scientific) and mounted. Until image acquisition via Leica SP8 lightning confocal microscopy (Wetzlar, Germany), slides were stored at -20°C.

**Table 2:** Antibodies used for either immunofluorescence (IF) or Western Blotting (WB)

| **Target** | **Antibody** | **Company** | **Dilution IF** | **Dilution WB** |
| --- | --- | --- | --- | --- |
| ACTA2 | ab124964 | Abcam<br>(Cambridge, UK) | 1:100 | - |
| ACTA2 | C6198 | Sigma-Aldrich (St. Louis, USA) | 1:250 | - |
| SOX2 | AF2018 | R&D Systems, Inc.<br>(Minneapolis, USA) | 1:100 | - |
| SOX9 | HPA001758 |  | 1:250 | - |
| pSMAD2/3 | 8828 | Cell Signaling<br>Technology, Inc.<br>(Danvers, USA) | - | 1:500 |
| pSMAD2 | 3101s | Cell Signaling<br>Technology, Inc.<br>(Danvers, USA) | 1:250 | - |
| Smad7 | ab216428 | Abcam<br>(Cambridge, UK) | 1:200 | 1:1000 |
| CD63 | ab59479 | Abcam<br>(Cambridge, UK) |  | 1:500 |
| TSG101 | 79644 | Cell Signaling Technology, Inc. (Danvers, USA) |  | 1:1000 |
| Flot1 | 610820 | BD Biosciences |  | 1:500 |
| CANX | ab22595 | Abcam (Cambridge, UK) |  | 1:1000 |
| IgG Alexa Fluor 488 Goat | Invitrogen/A11055 | Thermo Fisher Scientific (Waltham, USA) | 1:1000 | - |
| IgG Alexa Fluor 647 Rabbit | CST/4414S | Cell Signaling Technology, Inc. (Danvers, USA) | 1:1000 | - |
| IgG Alexa Fluor 555 Rabbit | CST/4413S | Cell Signaling Technology, Inc. (Danvers, USA) | 1:1000 | - |
| IgG HRP Rabbit | CST/7074 | Cell Signaling Technology, Inc. (Danvers, USA) | - | 1:1000 |
| IgG HRP Goat | Invitrogen/31402 | Thermo Fisher Scientific (Waltham, USA) | - | 1:1000 |
| IgG HRP Mouse | CST/7076S | Cell Signaling Technology, Inc. (Danvers, USA) | - | 1:1000 |

A minimum of 10 pictures from each sample was randomly taken for blinded quantification and analysis. ACTA2^+^ cell thickness was measured using ImageJ (1.8.0, National Institutes of Health) to calculate the corrected total cell fluorescence by subtracting the mean fluorescence of the background.

SOX2^+^ and SOX9^+^ nuclear staining was manually counted and normalized to the total number of DAPI^+^ cells.

### Bulk RNA-sequencing

Total RNA was isolated using RNeasy Plus Micro Kit (Qiagen) according to the manufacturer’s instructions. Three biological replicates per experimental group were chosen following an initial quality assessment step with bioinformatics tool FastQC. The library was prepared via Illumina Stranded Total RNA Prep Ligation with Ribo-Zero Plus kit. Sequencing of 100 million reads per sample was carried out on Illumina- NovaSeq2. DESeq2 and G-Profiler were employed for differential gene expression analysis.

### Antagomir studies

To inhibit miR-93-5p, we transfected AFSCs at 60% confluence with miR-93-5p inhibitor (25 μM) according to the manufacturer’s instructions (Qiagen Y104101029- DDA: power inhibitor rno-miR-93-5p; YI00199006-DDA: scrambled negative control). AFSCs were cultured and treated either with antagomir 93-5p or with scrambled control. The knock-out of miR-93-5p was validated within AFSCs as well as within their collected EVs via RT-qPCR for known targets of miR-93-5p such as Repulsive Guidance Molecule BMP Co-Receptor B (*Rgmb*) or Osteocalcin (*Ocn*).^43,44^ Hypoplastic lungs secondary to oligohydramnios were subsequently treated as described above and analyzed for Tgf-β signaling, lung branching morphogenesis and growth via RT-qPCR (*Tgfb1, Fgf10, Nrp1, Ctnnb1*) and histology (RAC, MLI)

### Western Blotting

Phosphatase and protease inhibitor (Roche) containing tissue extraction buffer (Thermo Fisher Scientific) was used to extract proteins from lung tissue samples. A bicinchoninic acid assay (Thermo Fisher Scientific) was used for quantification of proteins. Primary antibodies were incubated overnight at 4°C with subsequent 1h incubation with secondary antibodies (**Table 2**). The relative protein quantification was performed via Image Studio Light version 5.2 and normalized to the expression of ACTB.

### Statistical analysis

Statistical analysis was conducted by either using unpaired t (parametric) or Mann– Whitney U (non-parametric) test for comparison of two groups. One-way analysis of variance (ANOVA) (post hoc parametric Tukey comparison) or Kruskal-Wallis (post hoc Dunn’s nonparametric comparison) tests were performed for more than 2 groups. Normality of data was assessed according to Gaussian distribution assessed by D’Agostino-Pearson or Shapiro-Wilk normality test. P value < 0.05 was considered statistically significant. All analyses and graphs were created with GraphPad Prism software version 8.0.2.

## Results and Discussion

### Experimental oligohydramnios induces fetal pulmonary hypoplasia that is improved by administration of AFSC-EVs

First, we isolated AFSC-EVs using differential ultracentrifugation, as previously described (**Fig.1A**).^13,38^ This technique ensures high EV yield and is the most used isolation method to separate EVs for clinical applications in patients.^45^ From 10ml of AFSC conditioned medium, we extracted EVs with a typical double membrane morphology (**Fig.1B**), a concentration of 7.47x10^10^ and small size according to MISEV2023 (mean= 169±92.8nm; mode= 143nm; **Fig.1C**)^39^, and expressing canonical EV markers (CD63, TSG101, FLOT1) without evidence of cellular debris (lack of CANX protein expression; **Fig.1D**). It remains controversial whether AFSC-EV biological function varies according to vesicle size.^46^ In our previous study, we demonstrated that administration of small (140±5nm) but not large (363±17nm) AFSC-EVs improves lung development in fetal hypoplastic lungs secondary to CDH.^13^ On the other hand, other studies in models of renal fibrosis and skeletal muscle atrophy reported that also the administration of AFSC-EVs larger than 200 nm produced anti-fibrotic, angiogenic, and immunomodulatory effects.^47,48^ The size range and concentration of the AFSC-EVs employed in our experiments were in line with the mean size and yield of MSC-EVs used for the treatment of experimental bronchopulmonary dysplasia, an inflammatory neonatal lung disease that has shown promising response to MSC-EV regenerative properties.^49,50^

**Figure 1:**
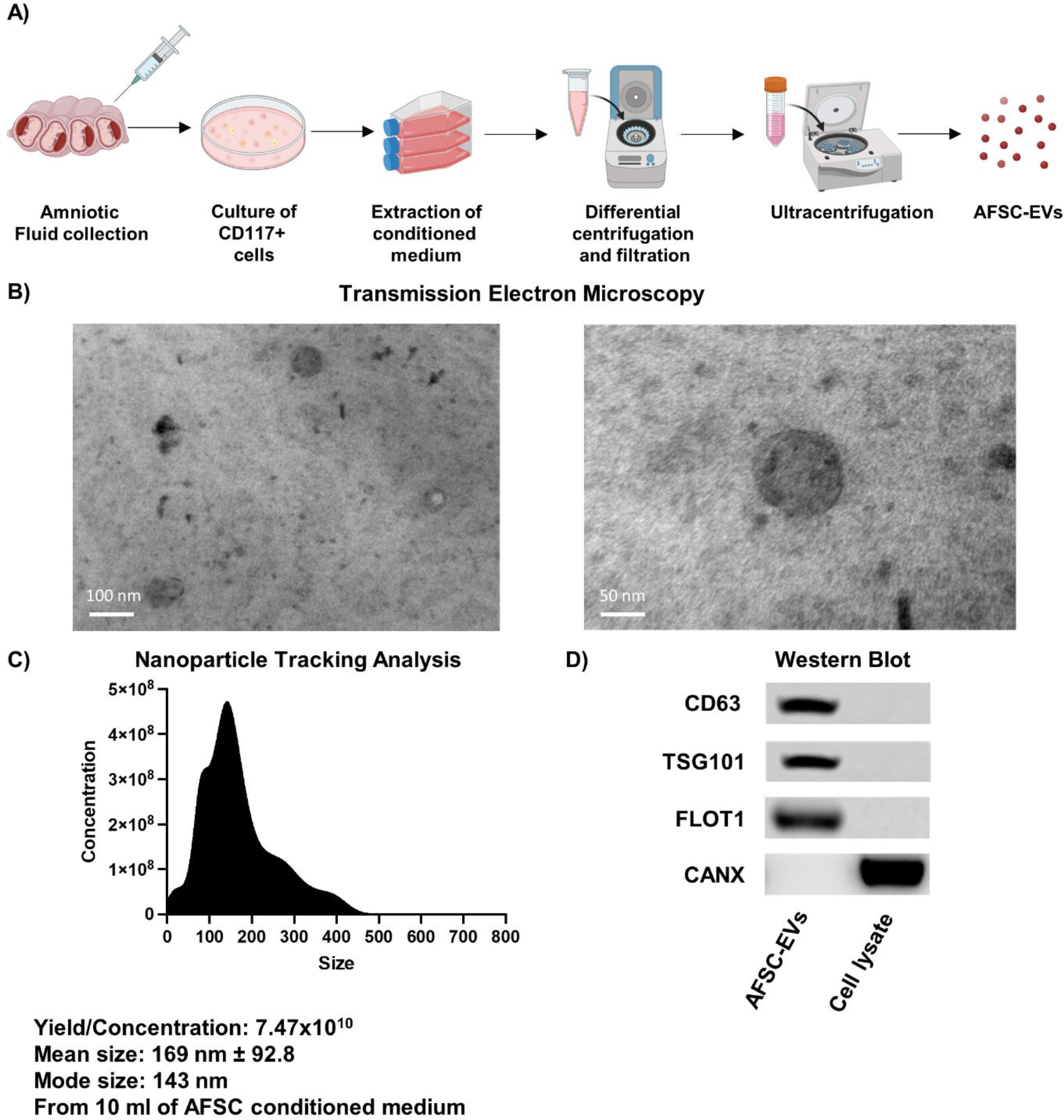
Isolation and characterization of extracellular vesicles (EVs) derived from amniotic fluid stem cells (AFSCs) **A)** Graphical overview of AFSC isolation and EV derivation from rat amniotic fluid. **B)** Representative image of the typical double membrane morphology of AFSC-EVs obtained via transmission electron microscopy (TEM) at low (left, scale: 100nm) and high (right, scale: 50nm) magnification **C)** Nanoparticle tracking analysis (NTA) distribution showing EV particle concentration relative to size (nm). **D)** Western blot analysis for canonical EV markers: cell differentiation marker 63 (CD63), tumor susceptibility gene 101 (TSG101), and flotillin-1 (Flot-1). Cell-specific marker calnexin- 1 (CANX) was used to exclude presence of cellular debris.

To induce pulmonary hypoplasia secondary to oligohydramnios, we employed a model based on amniotic fluid drainage in rat fetuses at embryonic day (E) 16.5 (late pseudoglandular stage of lung development) (**Fig.2A**), as previously described by another research group.^51,52^ In our experiments, fetuses with oligohydramnios (32 treated with medium alone and 18 treated with AFSC-EVs) had decreased lung-to-body weight ratio at E19.5 (canalicular stage) compared to 14 healthy age-matched controls (oligohydramnios: 3.1±0.4% vs. control: 3.8%±0.3%; p<0.0001) (**Fig.2B**). These data confirmed a decrease in bilateral lung weight, as previously described by Chen et al,^53^ and showed that lungs treated with medium alone or with AFSC-EVs had a similar weight at baseline (**Fig.2B**). As AFSC-EV regenerative effects to fetal hypoplastic lungs were mainly attributed to the delivery of their RNA cargo^13^, we first investigated the presence of AFSC-EV RNA signal in the lung and found it throughout the lung parenchyma (**Fig.2C**). We then compared controls to oligohydramnios+medium lungs and found that the latter had fewer airspaces and reduced expression of *Fgf10, Nrp1,* and *Ctnnb1* (**Fig.2D-E**). This picture of pulmonary hypoplasia is in line with that previously reported by others using the same model of oligohydramnios,^51^ as well as by us in the CDH pulmonary hypoplasia model.^13,33,54^ Treatment with AFSC-EVs improved lung growth as demonstrated by a higher fetal airspace density and rescued expression of *Fgf10*, *Nrp1,* and *Ctnnb1* back to control level (**Fig.2D-E**). This confirms our previous work demonstrating the effects of antenatal AFSC-EV administration in experimental fetal lung hypoplasia secondary to _CDH._13,33

**Figure 2:**
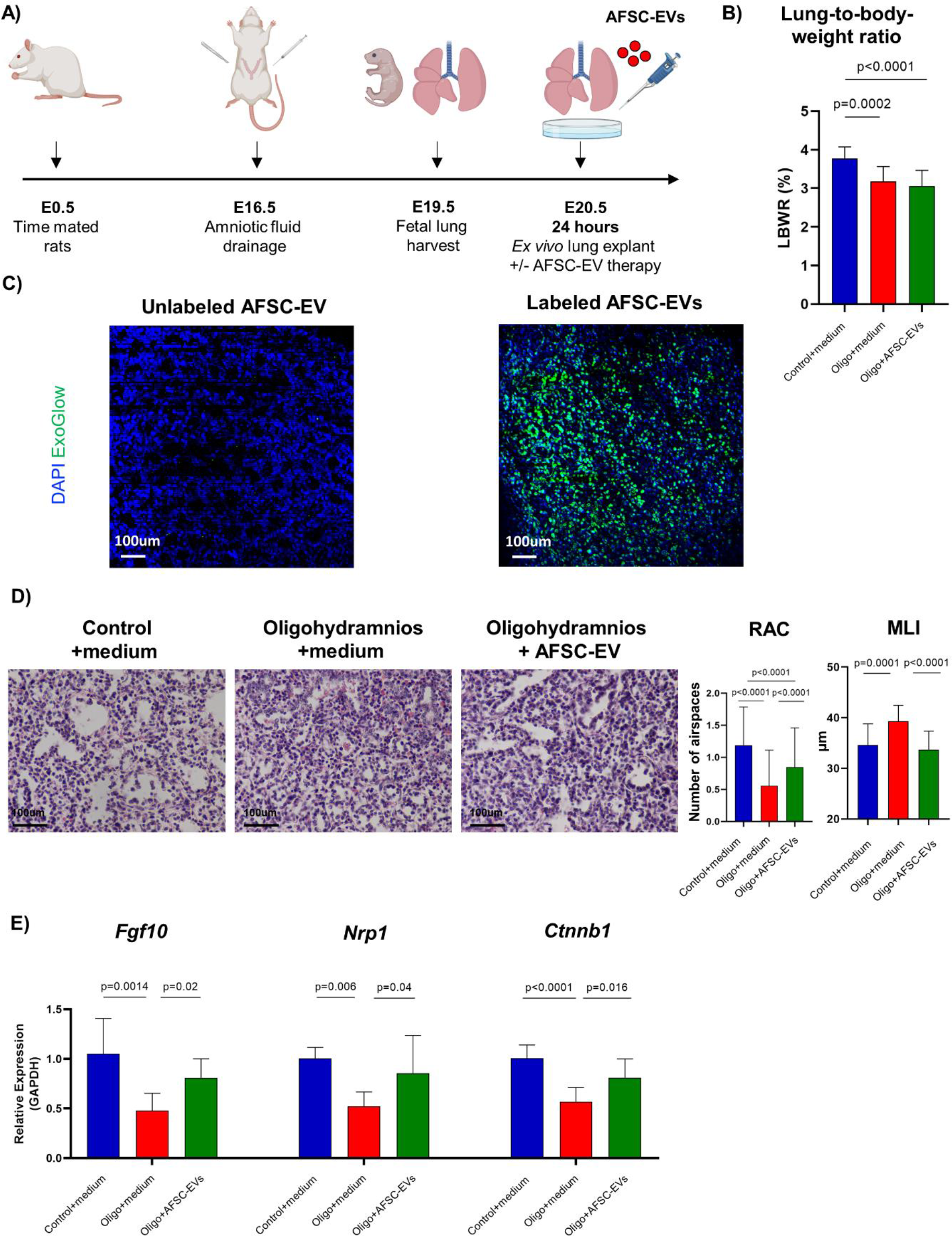
Experimental oligohydramnios induces fetal pulmonary hypoplasia with impaired fetal lung branching morphogenesis that is improved by administration of AFSC-EVs **A)** Graphical overview of the experimental timeline. **B)** Comparison of lung-to-body- weight ratio (LBWR, in %) between the three experimental groups at E19.5: control fetal rat lungs exposed to medium alone (Control+medium, blue), hypoplastic fetal rat lungs secondary to oligohydramnios treated with medium alone (Oligo+medium, red) or medium supplemented with AFSC-EVs (Oligo+AFSC-EVs, green) **C)** Uptake of AFSC-EV RNA cargo (green, Exo-GLOW™ labeled EVs) throughout the fetal lung parenchyma (blue, DAPI) at E19.5+1. **D)** Representative H&E staining from all three experimental groups with comparison of the radial airspace count (RAC, n) and the mean linear intercept (MLI, µm). Each group contains at least n=5 biological replicates and all technical replicates were included to determine statistical significance. **E)** Gene expression analysis in fetal lungs across all three experimental groups assessed via RT-qPCR for lung branching morphogenesis indicated by *Fgf10, Nrp1,* and *Ctnnb1*. Data is expressed by mean±standard deviation (SD). Each group contains at least n=6 biological replicates. Statistical significance was determined via one-way ANOVA (*Nrp1*, *Ctnnb1*) or Kruskal-Wallis test (LBWR, RAC, MLI, *Fgf10*) with post-hoc Tukey’s- or Dunn’s test.

### AFSC-EV treatment improves lung progenitor cell distribution in fetal hypoplastic lung explants secondary to oligohydramnios

As proximal and distal airway progenitor cells are known to orchestrate lung branching morphogenesis via interaction with airway smooth muscle cells that surround the developing airspaces, we studied the gene and protein expression and distribution of these cells in our oligohydramnios pulmonary hypoplasia model.^55^ We found that in comparison to controls, explants from rats with oligohydramnios treated with medium alone had significantly decreased expression of *Sox2* and *Sox9* (**Fig.3A**). To assess the airway spatial patterning, we quantified SOX2^+^, SOX9^+^, and ACTA2^+^ cells via immunofluorescence. We observed a significant decrease of SOX2^+^ and SOX9^+^ progenitor cells together with an increase of ACTA2^+^ smooth muscle cells in oligohydramnios fetal lung explants treated with medium alone in comparison to control fetal lung explants (**Fig.3B**). Qualitative analysis using immunofluorescence and triple staining for SOX2^+^, SOX9^+^, and ACTA2^+^ revealed severe disruption of the spatial airway patterning in oligohydramnios exposed fetal lung explants treated with medium alone (**Fig.3C**). The abnormal proximal-distal patterning and smooth muscle cell hypertrophy found in our model of oligohydramnios are in line with reports in other models of impaired development in rat and human fetal lungs.^55–57^ In our experiments, administration of AFSC-EVs to hypoplastic lung explants secondary to oligohydramnios resulted in a significant increase of *Sox2* and *Sox9* gene expression, as well as a rescue of SOX2^+^, SOX9^+^, and ACTA2^+^ protein expression levels (**Fig.3A- B**). Moreover, the distribution of SOX2^+^/SOX9^+^/ACTA2^+^ in the airways of AFSC-EV treated lungs resembled that of normal controls.

**Figure 3:**
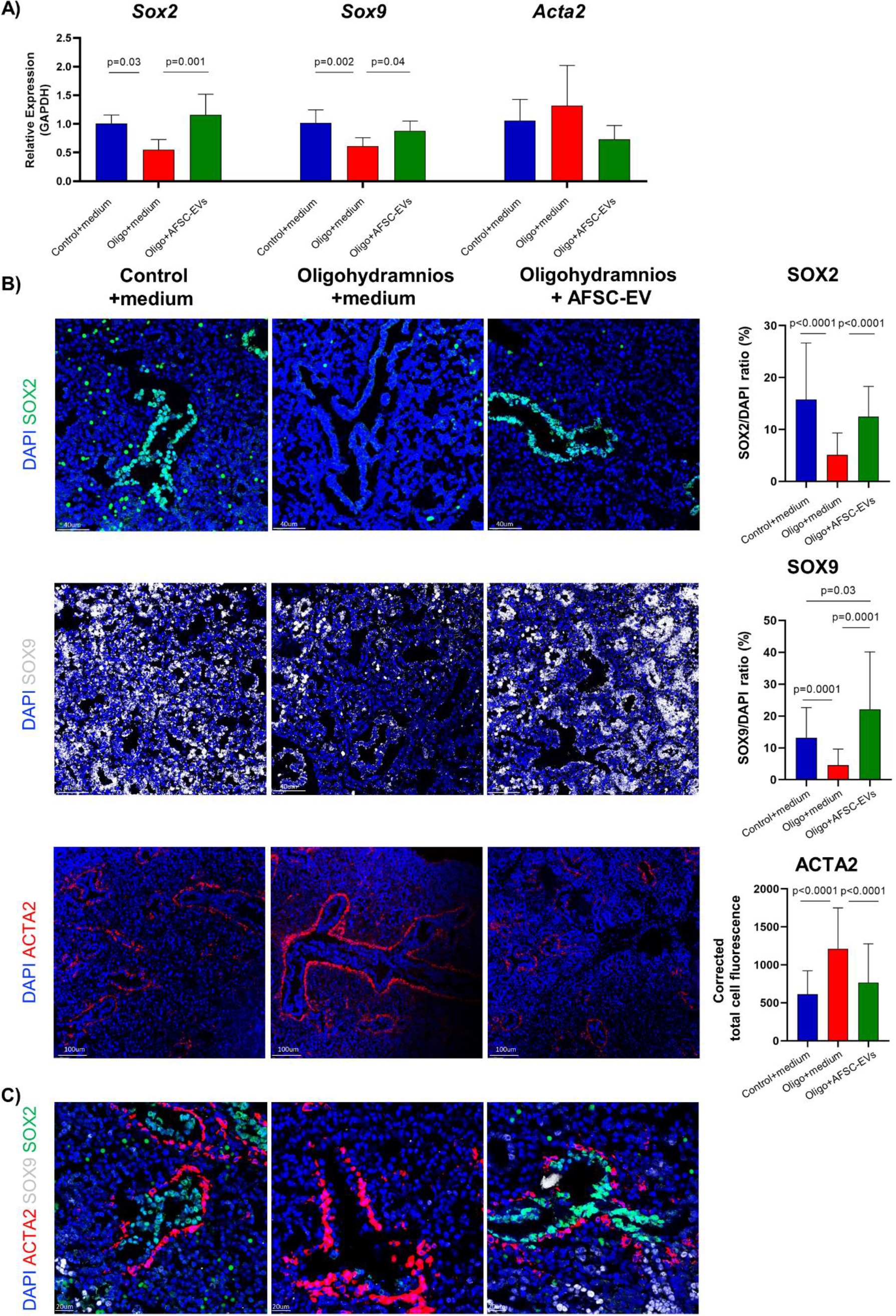
AFSC-EV treatment improves lung progenitor cell distribution in fetal hypoplastic lung explants secondary to oligohydramnios Gene level changes in fetal lungs assessed via RT-qPCR across three experimental groups: control fetal rat lungs exposed to medium alone (Control+medium, blue), hypoplastic fetal rat lungs secondary to oligohydramnios treated with medium alone (Oligo+medium, red) or medium supplemented with AFSC-EVs (Oligo+AFSC-EVs, green) for lung progenitor cell markers (*Sox2* and *Sox9*) and smooth muscle cells (*Acta2*) at E19.5+1. Each group contains at least n=7 biological replicates. **B-C)** Protein expression changes and representative images of single or triple staining for SOX2^+^, SOX9^+^, ACTA2^+^ and DAPI^+^ across experimental groups assessed via immunofluorescence. At least n=5 biological replicates were used per group and all technical replicates have been used for statistical analysis. Statistical significance was determined via one-way ANOVA (*Sox2*, *Sox9*, *Acta2*) or Kruskal Wallis test (SOX2, SOX9, ACTA2) with post-hoc Tukey’s or Dunn’s test and is presented as mean±SD.

### Induction of oligohydramnios leads to transcriptomic alterations in fetal lung explants that are partly improved via AFSC-EV administration

To understand the effects of AFSC-EVs on lung branching morphogenesis, we conducted RNA-sequencing of the lung explants from all three conditions. Differential gene expression analysis between oligohydramnios+medium lung explants and controls revealed 118 significantly up- and 80 significantly down-regulated genes, which annotated to pathways related to multicellular organismal processes, extracellular space, cell surface, adhesion, and periphery (**Fig.4A**). These biological functions are key for the organization of the developing airways. On the one hand, cell surface and adhesion molecules guide epithelial cells entering the adjacent structures to enable lung branching morphogenesis and spatial progenitor cell distribution to provide a protective barrier.^58^ On the other hand, the extracellular matrix provides a support to lung cells and is essential for biophysical and biochemical signaling between them.^59,60^ In our analysis, we also found that compared to oligohydramnios+medium lung explants, oligohydramnios+AFSC-EV lung explants had 14 downregulated genes that are involved in biological processes related to extracellular space regulation, tissue development, and cell differentiation (**Fig.4B**). This confirms previous reports showing that fetal hypoplastic lungs have an epithelial-mesenchymal imbalance with dysregulated mesenchymal and epithelial cell distribution and differentiation.^33,61,62^ Moreover, we previously proved that AFSC-EV administration improves lung tissue development by promoting epithelial and mesenchymal cell differentiation.^33^ In our current analysis, of the 14 downregulated genes in oligohydramnios+AFSV-EV lung explants, 11 were upregulated in oligohydramnios+medium lung explants compared to controls (**Fig.4C**).

**Figure 4:**
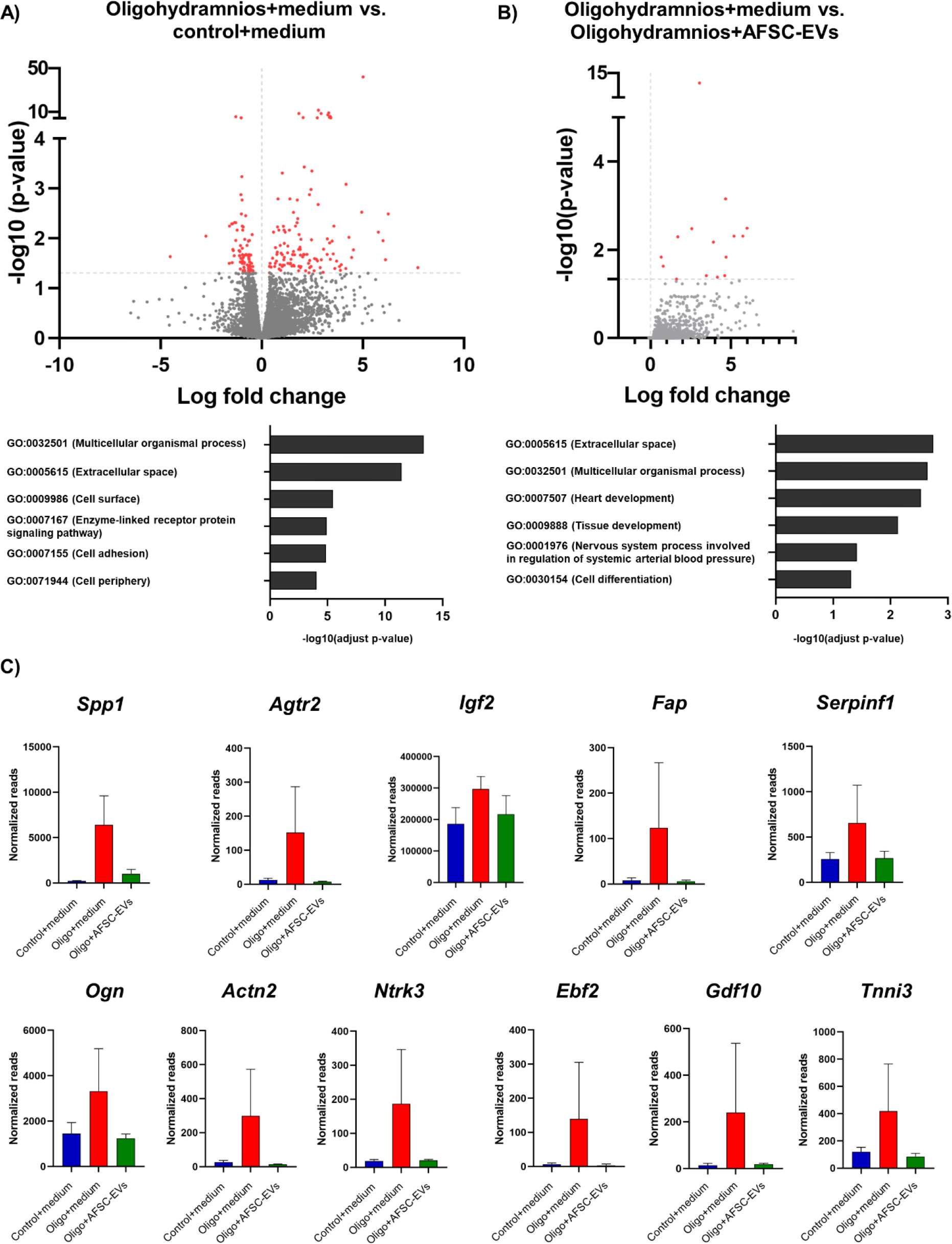
RNA-sequencing analysis of hypoplastic lungs secondary to oligohydramnios reveals transcriptomic alterations that are partly improved via AFSC-EV administration. **A)** Volcano plot illustrating log fold-change plotted against adjusted p-value (-log base 10) of differentially expressed genes comparing hypoplastic fetal rat lungs secondary to oligohydramnios (n=3 biological replicates) and control fetal rat lungs (n=3 biological replicates), both exposed to medium alone. Top 6 pathways determined via gene set enrichment analysis of GO terms (g:Profiler) and expressed as the adjusted p-value (- log base 10). **B)** Volcano plot illustrating log fold-change plotted against adjusted p-value (-log base 10) of differentially expressed genes comparing hypoplastic fetal rat lungs secondary to oligohydramnios treated with medium alone (n=3 biological replicates) or AFSC-EVs (n=3 biological replicates). Top 6 pathways determined via gene set enrichment analysis of GO terms (G-profiler) and expressed as the adjusted p-value (-log base 10). **C)** Number of normalized reads of 11 differentially expressed genes that are upregulated in oligohydramnios exposed fetal lungs (Oligo+medium, red) in comparison to control fetal lungs (Control+medium, blue) that were downregulated by AFSC-EV administration (Oligo+AFSC-EVs, green).

The majority of these 11 genes have been reported to be implicated in the Tissue Growth Factor beta (TGF-β) signaling pathway (**Table 3**). This pathway is known to play a crucial role throughout embryogenesis and organogenesis from the very early stages of development.^63^ In the developing lung, TGF-β signaling is important for branching morphogenesis and alveolarization, and promotes epithelial-mesenchymal cell interactions.^64^ This has been confirmed by studies employing TGF-β knock-out models that resulted in lethal pulmonary hypoplasia at birth and knock-out of its downstream targets SMAD3 that led to reduced pulmonary alveolarization.^64–67^ Moreover, TGF-β signaling is implicated in mechano-transduction and its expression has been reported to be altered in experimental oligohydramnios.^52,68^ Specifically, Chen et al reported that in a model of oligohydramnios, fetal rats had hypoplastic lungs with decreased gene and protein expression of TGF-β1.^52^ More recently, Callaway et al established a mouse model of oligohydramnios, where they showed that TGF-β controls lung epithelial cell plasticity.^68^ These studies and our current data support the knowledge that fetal lungs require TGF-β signaling for normal lung development and that oligohydramnios lungs have dysregulation of this pathway.

**Table 3:**
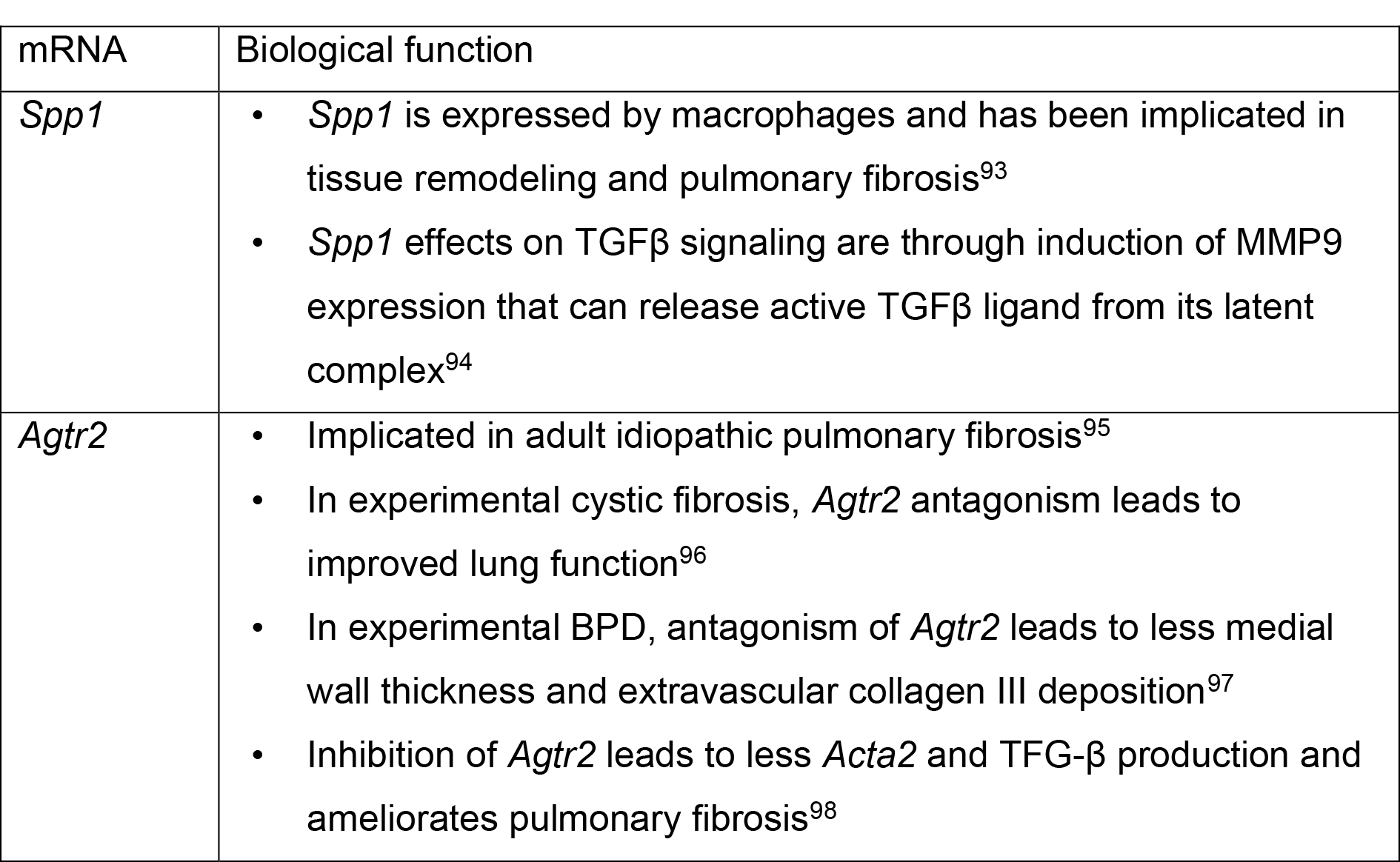
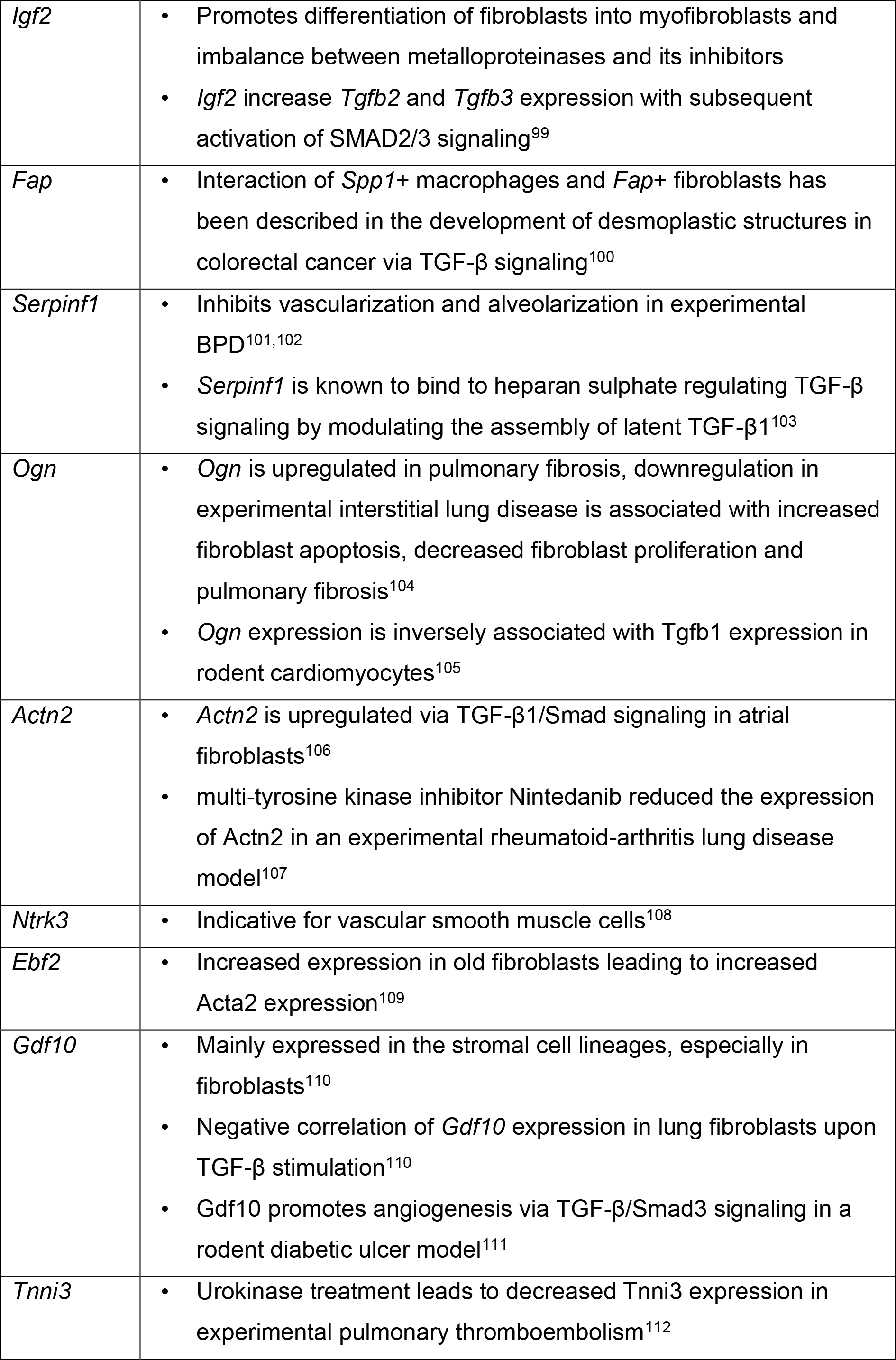
Differentially expressed genes across all three experimental groups and their biological function.

### AFSC-EV administration restores the TGF-β signaling pathway in hypoplastic lungs of rat fetuses with oligohydramnios

When we investigated the expression of factors involved in the TGF-β signaling pathway, we found that oligohydramnios+medium lung explants had decreased levels of *Tgfb1* and its receptors *Tgfbr1* and *Tgfbr2* in comparison to control fetal lung explants (**Fig.5A**). Administration of AFSC-EVs rescued the gene expression of *Tgfb1* back to control level (**Fig.5A**). We also found that phosphorylated SMAD2 and 3 (pSMAD2/3) were downregulated in oligohydramnios+medium fetal lung explants in comparison to healthy controls and that administration of AFSC-EVs rescued pSMAD2/3 expression (**Fig.5B**). Phosphorylation of SMAD2/3 causes their translocation and accumulation in the nucleus, where they directly induce transcription of genes that regulate cell proliferation, differentiation, and migration.^69,70^ We also investigated the expression of SMAD 7, a protein that blocks the cascade and induces signal termination.^69^ In our experiments, the expression of SMAD 7 was rescued to normal levels by AFSC-EV administration (**Fig.5C**). These data are in line with previous reports showing that EVs derived from different sources of stem cells target SMAD proteins in various disease models.^71,72^

**Figure 5:**
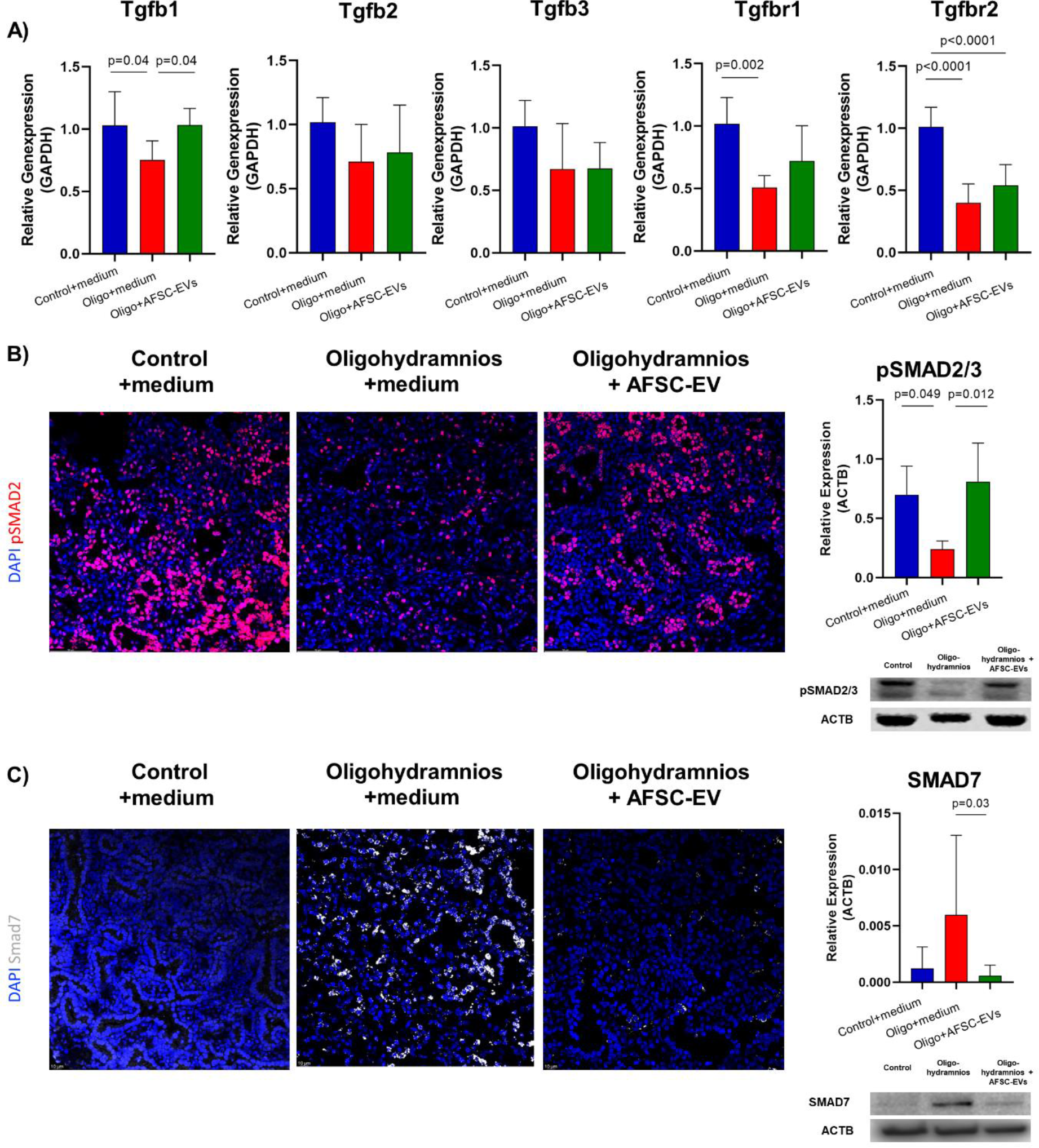
AFSC-EV administration restores the TGF-β signaling pathway in hypoplastic lungs of rat fetuses with oligohydramnios. **A)** Gene expression changes across experimental groups: control fetal rat lungs exposed to medium alone (Control+medium, blue), hypoplastic fetal rat lungs secondary to oligohydramnios treated with medium alone (Oligo+medium, red) or medium supplemented with AFSC-EVs (Oligo+AFSC-EVs, green) assessed via RT-qPCR for the three TGF-β isoforms (*Tgfb1*, *Tgfb2*, *Tgfb3*) and their 2 receptors (*Tgfbr1*, *Tgfbr2*) at E19.5+1. Each group contains at least n=6 biological replicates except of *Tgfb3* (at least n=3 biological replicates). **B-C)** Representative immunofluorescence staining and protein level changes for pSMAD2/3 and SMAD7 across all three experimental groups at E19.5+1 assessed via Western Blotting. Each group contains at least n=4 biological replicates. Statistical significance was determined via one-way ANOVA (*Tgfb1*, *Tgfb3*, *Tgfbr2*, pSMAD2/3) or Kruskal Wallis test (*Tgfb2*, *Tgfbr1*, SMAD7) with post-hoc Tukey’s or Dunn’s test and is presented as mean±SD.

### MiR-93-5p is critically involved in rescuing impaired lung branching morphogenesis and lung growth secondary to oligohydramnios

As we previously reported that AFSC-EV effects on another model of fetal pulmonary hypoplasia is at least in part mediated by the delivery of their miRNA cargo^13^, we first interrogated miRwalk for miRNAs that control the 14 genes that our sequencing analysis found downregulated in oligohydramnios+AFSC-EV lung explants (**Fig.6A**). We then cross-examined this list of miRNAs with the most abundant miRNAs present in the AFSC-EV cargo that were identified with AFSC-EV small RNA sequencing (**Fig.6A**).^13^ The network that we built revealed several miRNAs that are important for lung development, such as the miR17∼92 cluster and its paralogs. In particular, we found that AFSC-EVs are highly enriched in miR-93-5p, a regulator of the TGF-β signaling pathway that is predicted to modulate the expression of 12 of the 14 downregulated genes.^73–75^ Moreover, miR-93-5p has been reported to modulate TGF- β signaling through the negative regulator SMAD 7, which we found downregulated in oligohydramnios+AFSC-EV explants compared to oligohydramnios+medium lung explants.^69,76–79^ Lastly, miRWalk database reported that miR-93-5p is predicted to bind all the factors implicated in branching morphogenesis and TGF-β signaling that we found downregulated in our experiments (*Fgf10*, *Nrp1*, *Ctnnb1*, *Sox2*, *Sox9*, *Tgfb1*, *Tgfbr1*, *Tgfbr2*, *Smad2*, and *Smad3;* **Fig.6A**). Given this potential pivotal effect of miR- 93-5p, we hypothesized that antagonizing its expression in AFSC-EVs would result in a loss of AFSC-EV beneficial effect on branching morphogenesis. To conduct inhibition studies, we transfected parental AFSCs with a miR-93-5p antagomir or with a control scrambled sequence (**Fig.6B**). We first confirmed the knock-down of miR-93-5p in AFSCs and their derived EVs by the upregulation of validated miR-93-5p targets, such as *Rgmb* and *Ocn* (**SFig.1**). When we treated oligohydramnios hypoplastic lungs with AFSC-EVs transfected via antagomir 93-5p, we observed a significant downregulation of *Tgfb1* gene expression compared to explants treated with wild-type AFSC-EVs (**Fig.6C**). Similarly, oligohydramnios hypoplastic lungs treated with antagomir 93-5p transfected AFSC-EVs had significantly downregulated expression of *Fgf10*, *Nrp1* and *Ctnnb1* compared to explants treated with wild-type AFSC-EVs (**Fig.6C**). We also confirmed that oligohydramnios hypoplastic lungs treated with transfected antagomir 93-5p AFSC-EVs had impaired branching morphogenesis compared to explants treated with wild-type AFSC-EVs or the scrambled sequence control (**Fig.6D**). In summary, our data suggest that miR-93-5p inhibits SMAD7 and thus enhances TGF- β signaling via SMAD2 and SMAD3 phosphorylation (**Fig.6E**).

**Figure 6:**
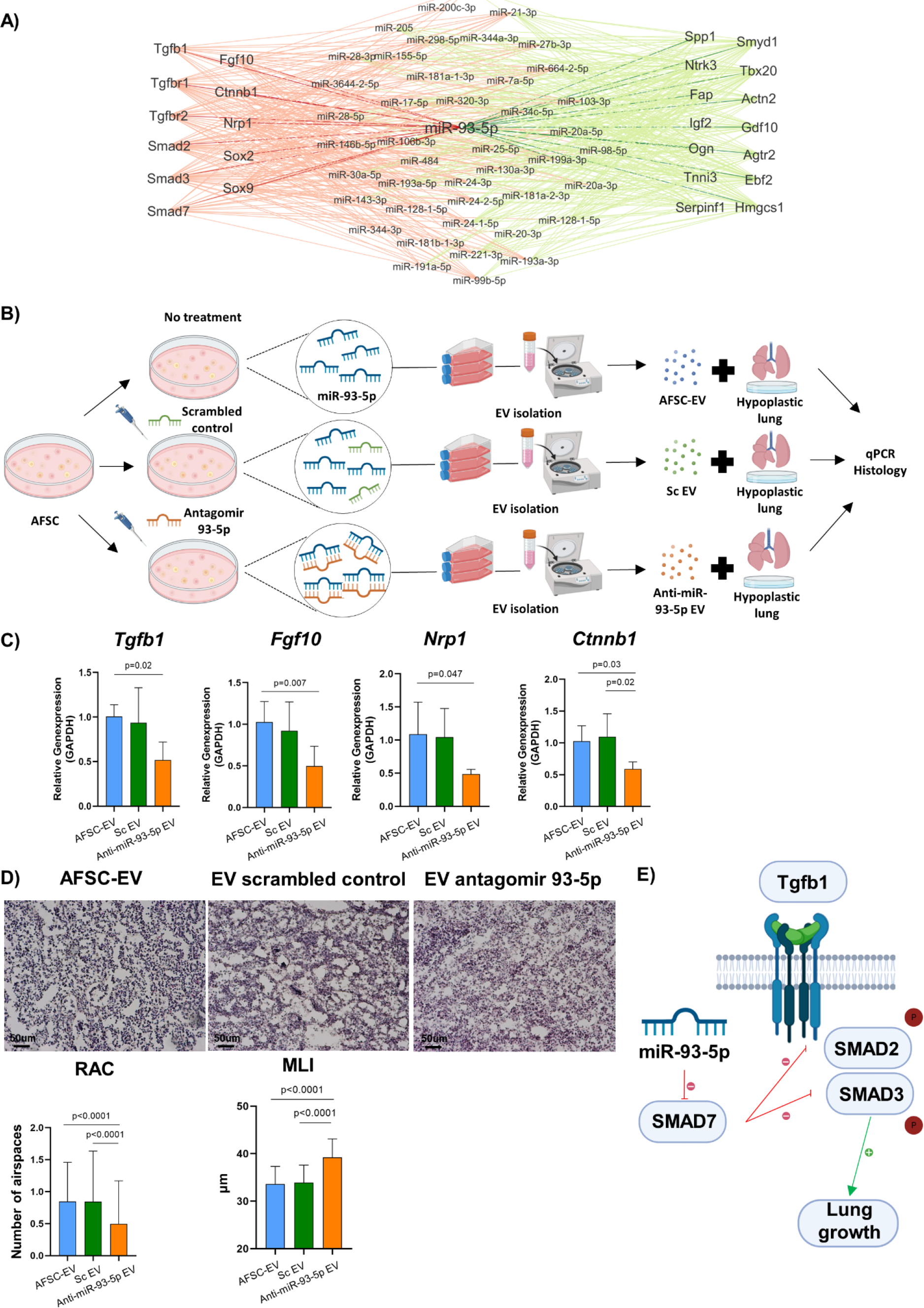
MiR-93-5p is critically involved in rescuing impaired lung branching morphogenesis and lung growth secondary to oligohydramnios. Predicted interaction network of miRNAs contained in the AFSC-EV cargo and genes involved in lung branching morphogenesis, organ growth, and TGF-β signaling pathways. **B)** Experimental timeline of antagomir studies to knock-down the expression of miR-93-5p within AFSC-EVs. **C)** Gene expression of *Tgfb1, Fgf10, Nrp1* and *Ctnnb1* within the experimental groups: hypoplastic fetal lung explants secondary to oligohydramnios and treated either with wild-type (AFSC-EV, blue), scrambled control treated (ScEV, green) or antagomir 93-5p treated (Anti-miR-93-5p EV, orange) EVs. Each group contains at least n=4 biological replicates **D)** Representative H&E staining from all experimental groups with comparison of the mean linear intercept (MLI, µm) and radial airspace count (RAC, n) at E19.5+1. Each group contains at least n=3 biological replicates and all technical replicates were included to determine statistical significance (technical replicates). **E)** Schematic overview of improved TGF-β/SMAD signaling via miR-93-5p inhibition of SMAD7. Statistical significance was determined via one-way ANOVA (*Tgfb1*, *Fgf10, Nrp1, Ctnnb1,* MLI) or Kruskal Wallis test (RAC) with post-hoc Tukey’s or Dunn’s test and is presented as mean±SD.

Overall, our study shows promise for an EV-based therapy to promote lung development in an experimental model of oligohydramnios. EV-based therapies are emerging as a potential new treatment strategy for various conditions, as evidenced by several ongoing clinical trials testing EVs in patients with neoplastic diseases, graft versus host disease, Covid-19, Alzheimer’s disease, Meniere’s disease, and osteoarthritis.^45,80–88^ Most of these studies were safety and feasibility phase I/II trials and have shown no undesired effects in the recipients.^45^

We acknowledge that our current study has some limitations that need to be addressed before translating this promising EV-based therapy into clinical practice. The findings herein presented were obtained in an *ex-vivo* model of oligohydramnios. Further studies will address the application of AFSC-EVs in an *in-vivo* model, where route of administration, EV dose and frequency will also be tested. Moreover, the *in-vivo* model will allow us to conduct EV tracking studies and investigate where AFSC-EVs home and whether their uptake by other organs beyond the fetal lung could be associated with negative undesired side effects. In our previous studies using AFSC-EVs for the treatment of pulmonary hypoplasia secondary to CDH, we showed that intra-amniotically injected AFSC-EVs are present not only in the lung but also in the liver, kidney, intestine, and brain.^89^ Interestingly, localization of AFSC-EVs in the fetal brain of rats with CDH was associated with reduced brain tissue inflammation, evidenced by a decreased activation of microglia.^90^ Lastly, testing administration of GMP-grade AFSC-EVs in a large animal model of oligohydramnios, such as the fetal lamb model, will allow us to test their pharmacokinetic and pharmacodynamic properties, as well as the safety for the mother, as EVs are able to cross the placental barrier.^91^ Increasing clinical usage of EVs together with our experimental data warrant further investigations on stem cell derived EVs as a potential novel regenerative therapy.

## Conclusions

This study has shown that AFSC-EVs hold regenerative potential in oligohydramnios induced fetal pulmonary hypoplasia. Antenatal AFSC-EV administration improved features of lung development at least in part via the release of miR-93-5p. MiR-93-5p modulates TGF-β signaling, which is known to be implicated in various neonatal diseases such as bronchopulmonary dysplasia and CDH.^16,92^ Experimental stem-cell based therapies for pulmonary diseases are emerging and this study implies the potential use of AFSC-EVs to promote normal lung development in fetal hypoplastic lungs secondary to oligohydramnios.

## Acknowledgements

We are thankful for the assistance of the Imaging Facility, the Nanoscale Biomedical Imaging Facility, and the Lab Animal Services at the Hospital for Sick Children (SickKids), Toronto. The graphical schemes were designed using BioRender.com. Funding: this project was supported by Canadian Institutes of Health Research (CIHR) -CIHR Project Grant (175300 and 497959, AZ), CIHR Fellowship (187855, RLF; 176535, KK), and the German Research Foundation (#466815475, FD)

## Data availability statement

Bulk-RNA Sequencing data is available on GEO GSE247978.

---

**Supplementary figure 1:**
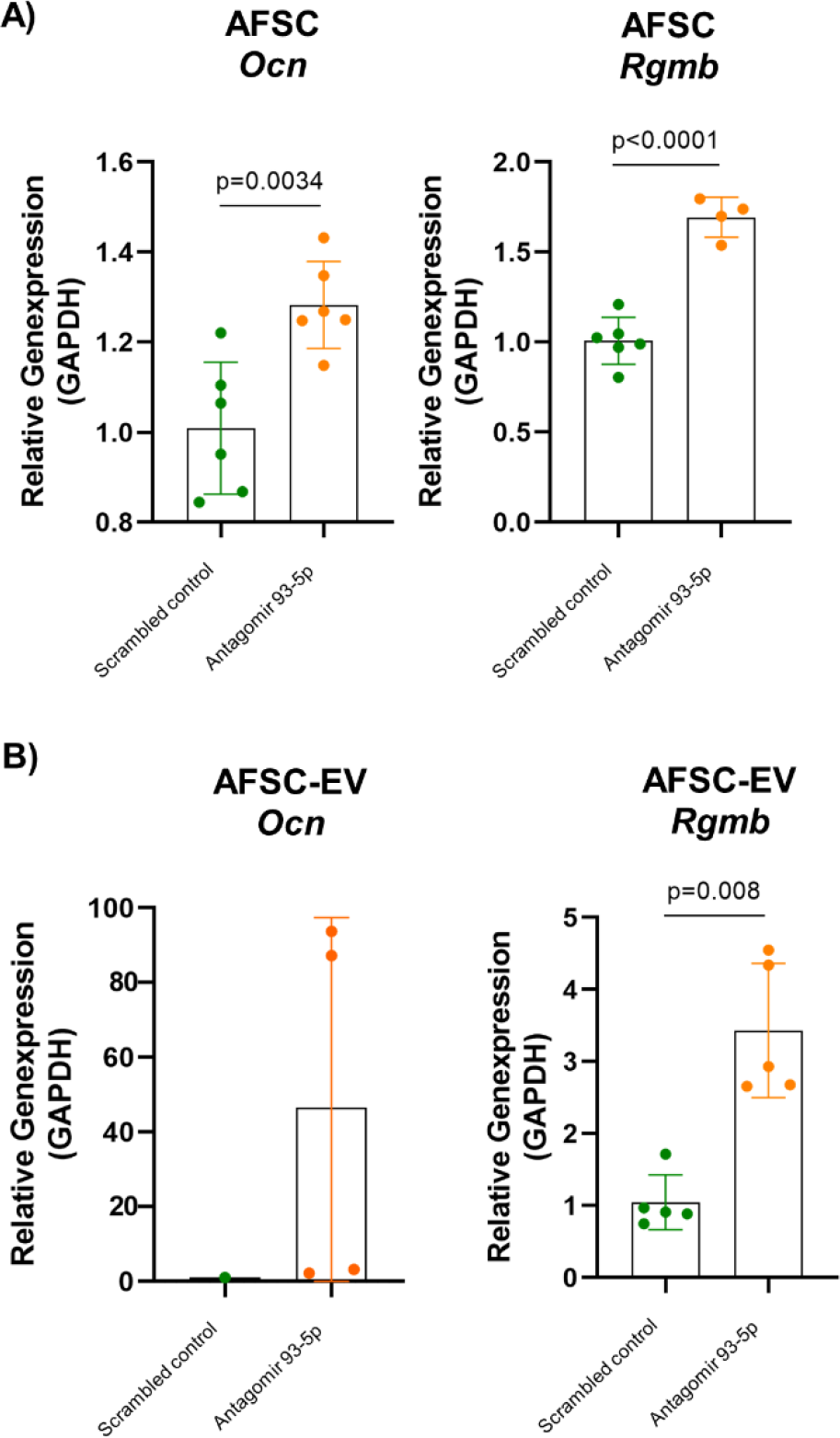
Validation of antagomir 93-5p transfection in AFSCs and AFSC-EVs. Gene expression changes assessed via RT-qPCR between scrambled control (green) and antagomir 93-5p (orange) transfected **A)** AFSCs and **B)** AFSC-EV for targets of miR-93-5p: Osteocalcin (*Ocn*) and repulsive guidance molecule BMP co-receptor b (*Rgmb*). Statistical significance was determined via unpaired t- (*Ocn*, *Rgmb* in AFSCs) or Mann Whitney U-test (*Rgmb* in AFSC-EVs) and is presented as mean±SD.

