## Supplementary figure 1 for "Amniotic fluid stem cell extracellular vesicles promote lung development via TGF-beta modulation in a fetal rat model of oligohydramnios"

**Supplementary figure 1:**

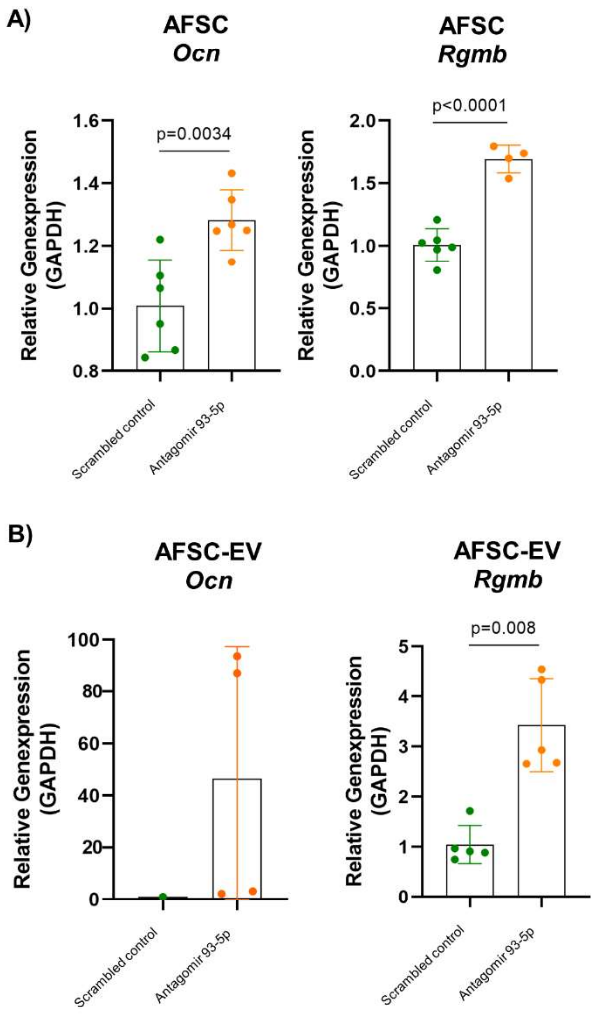


**Supplementary figure 1: Validation of antagomir 93-5p transfection in AFSCs and AFSC-EVs**

Gene expression changes assessed via RT-qPCR between scrambled control (green) and antagomir 93-5p (orange) transfected **A)** AFSCs and **B)** AFSC-EV for targets of miR-93-5p: Osteocalcin (*Ocn*) and repulsive guidance molecule BMP co-receptor b (*Rgmb*). Statistical significance was determined via unpaired t- (*Ocn*, *Rgmb* in AFSCs) or Mann Whitney U-test (*Rgmb* in AFSC-EVs) and is presented as mean±SD.
